# Decreased overall neuronal activity in a rodent model of impaired consciousness during absence seizures

**DOI:** 10.1101/2021.04.20.440390

**Authors:** Cian McCafferty, Benjamin Gruenbaum, Renee Tung, Jing-Jing Li, Peter Salvino, Peter Vincent, Zachary Kratochvil, Jun Hwan Ryu, Aya Khalaf, Kohl Swift, Rashid Akbari, Wasif Islam, Prince Antwi, Emily Ann Johnson, Petr Vitkovskiy, James Sampognaro, Isaac Freedman, Adam Kundishora, Basavaraju G. Sanganahalli, Peter Herman, Fahmeed Hyder, Vincenzo Crunelli, Antoine Depaulis, Hal Blumenfeld

**Affiliations:** University College Cork; Yale University; Cardiff University; Grenoble Institut des Neurosciences

**Keywords:** absence seizure, behavior, epilepsy, thalamus, consciousness, spike-wave-discharge, somatosensory cortex, ensemble recordings, functional magnetic resonance imaging

## Abstract

Absence seizures are characterized by a brief behavioural impairment including apparent loss of consciousness. Neuronal mechanisms determining the behavioural impairment of absence seizures remain unknown, and their elucidation might highlight therapeutic options for reducing seizure severity. However, recent studies have questioned the similarity of animal spike-wave-discharges (SWD) to human absence seizures both behaviourally and neuronally. Here, we report that Genetic Absence Epilepsy Rats from Strasbourg recapitulate the decreased neuroimaging signals and loss of consciousness characteristic of human absence seizures. Overall neuronal firing is decreased but rhythmic in the somatosensory cortex and thalamus during these seizures. Interestingly, individual neurons in both regions tend to consistently express one of four distinct patterns of seizure-associated activity. These patterns differ in firing rate dynamics and in rhythmicity during seizure. One group of neurons showed a transient initial peak in firing at SWD onset, accounting for the brief initial increase in overall neuronal firing seen in cortex and thalamus. The largest group of neurons in both cortex and thalamus showed sustained decreases in firing during SWD. Other neurons showed either sustained increases or no change in firing. These findings suggest that certain classes of cortical and thalamic neurons may be particularly responsible for the paroxysmal oscillations and consequent loss of consciousness in absence epilepsy.

## Introduction

Absence epilepsy is a condition defined by recurrent non-convulsive seizures involving brief (~5-30 second) behavioural arrest and electrographic spike-wave-discharges (SWD) (Blumenfeld, 2005). These SWDs are reflective of generalized paroxysmal brain activity, with oscillations particularly prevalent in the cerebral cortex and thalamus (McCormick and Contreras, 2001). Recent studies in multiple animal models of SWDs have advanced understanding of the precise contributions of neuronal populations to these paroxysms. Key features identified include a subset of hyper-excitable deep somatosensory cortical neurons that appear to initiate SWDs (Polack et al., 2007), a leading role of somatosensory cortex in the initial phase of the oscillation (Meeren et al., 2002), and an excitatory drive from cortical (but not thalamocortical) to reticular thalamic neurons throughout an SWD (McCafferty et al., 2018). Furthermore, there is some diversity in cortical and thalamocortical neuronal activity both within and between seizures (McCafferty et al., 2018; Meyer et al., 2018). This diversity raises the possibility that certain components of neuronal activity during seizure may be of particular significance in determining the behavioural features of absence. The specifics of seizure-associated neuronal activity can be directly investigated in animal models of absence epilepsy.

There have, however, been recent suggestions that animal SWDs, particularly in rodents, do not usefully resemble those that are part of absence seizures in humans despite their established pharmacological similarities (Crunelli et al., 2020). Rats can vary the duration of their SWDs in order to obtain rewards (Taylor et al., 2017) and SWDs are present to some extent in wild-caught rats (Taylor et al., 2019). Neither these observations nor previous studies (Vergnes et al., 1991) have addressed the fundamental question of whether rodents lose consciousness, and thus the ability to carry out directed behaviours, during SWDs. This may be because of the relationship between absence seizures and arousal level, in both humans and animal models: seizures are more likely to occur in a state of relaxed wakefulness and tend to be entirely suppressed during active engagement in a task (Horita et al., 1991; Zarowski et al., 2011). As such, conventional tests of consciousness, perception and responsiveness cannot be presented during animal SWDs. Another limitation of previous rodent absence models has been the finding of fMRI increases during SWD, whereas human absence seizures show mainly fMRI decreases in widespread cortical networks (Bai et al., 2010; Berman et al., 2010; Guo et al., 2016).

We sought first to replicate human neuroimaging findings in absence epilepsy by developing a novel model using awake, restraint-habituated GAERS, thereby eliminating the potential confound of anesthetic agents used in prior fMRI studies. In addition, in this study we devised modified paradigms to investigate behaviour and consciousness in GAERS during SWDs. These paradigms, analogous to those previously used in human studies, demonstrated a broad impairment of desirable behaviour during the SWDs. During a small proportion of electrographically distinct events, certain behaviours were spared – an observation that has also been made in humans (Guo et al., 2016). Using ensemble electrophysiological recordings with spike sorting, we then showed that overall neuronal activity in both cortex and thalamus is decreased during these behaviour-impairing SWDs. We also noted that this overall decrease is consistently driven by a subset of excitatory neurons in both regions, with the remaining neurons falling into one of three other seizure-associated activity patterns.

## Results

### Neuroimaging signal decreases during GAERS SWD resemble human absence seizures

Human research in children with typical absence seizures has shown prominent fMRI decreases in most cortical regions during generalized SWD (Bai et al., 2010; Berman et al., 2010). However, previous work in rodent models of absence epilepsy, mostly done under anaesthesia, has yielded variable results, with most studies showing cortical fMRI increases during SWD (Tenney et al., 2003, 2004; Nersesyan et al., 2004; David et al., 2008; Mishra et al., 2011). It was proposed that fMRI increases in rodent models were produced by anaesthetic agents not used in human studies; this proposal was further supported by observation of fMRI increases in an anesthetized ferret model, despite that fact that the SWD had similar frequency to humans and the thalamocortical anatomy was more similar to humans than in rodent models (Youngblood et al., 2015). We therefore sought to study neuroimaging signals in a rodent absence epilepsy model without anaesthetic agents. This was accomplished by gradual habituation of the GAERS model to the ambient noise and cloth body restraint used in the neuroimaging environment. In this awake model of spontaneous undrugged SWD, we found that both fMRI and laser Doppler flowmetry signals from the cortex showed predominantly decreased activity. The unanaesthetised GAERS model therefore shows cortical neuroimaging signal decreases during SWD resembling human absence seizures.

### Impaired behaviour during GAERS SWD resembles human absence seizures

We investigated the behavioural consequences of GAERS SWDs using two different paradigms, designed to be analogous to the repetitive tapping task (RTT) and continuous performance task (CPT) previously employed to study human behaviour during absence seizures (Guo et al., 2016). It has been challenging in previous rodent absence model work to demonstrate impaired behavior during SWD, at least in part because behavioral tasks increase arousal which tends to supress or interrupt SWD, preventing testing of behavior during SWD without interruption (van Luijtelaar et al., 1991; Smyk et al., 2011). Therefore, in our studies both tasks were modified to maintain a level of engagement/arousal compatible with uninterrupted seizure expression. In one task, designed to echo the human CPT, we conditioned GAERS to respond within 10 seconds to an 80 dB pure auditory tone at 8 kHz, presented at 60 s intervals, in order to receive a sucrose water reward. The inter-stimulus interval was gradually increased to an average of 180 s (or the automated detection of a SWD based on EEG amplitude) and the intensity decreased to 45 dB. During baseline periods, animals had a 88.2 ± 2.8% response rate after conditioning (4482 total stimuli in 14 rats) while during SWDs the response rate was decreased (p = 0.00012, Wilcoxon signed rank test) to 0.4 ± 0.3% (156 total stimuli). This behaviour appeared to be restored immediately post-SWD (p = 0.2163 compared to pre-seizure, Wilcoxon signed rank test), with response rates recovering to 78.2 ± 6.8% (330 post-ictal stimuli) (Fig. 1C).

**Figure 1.**
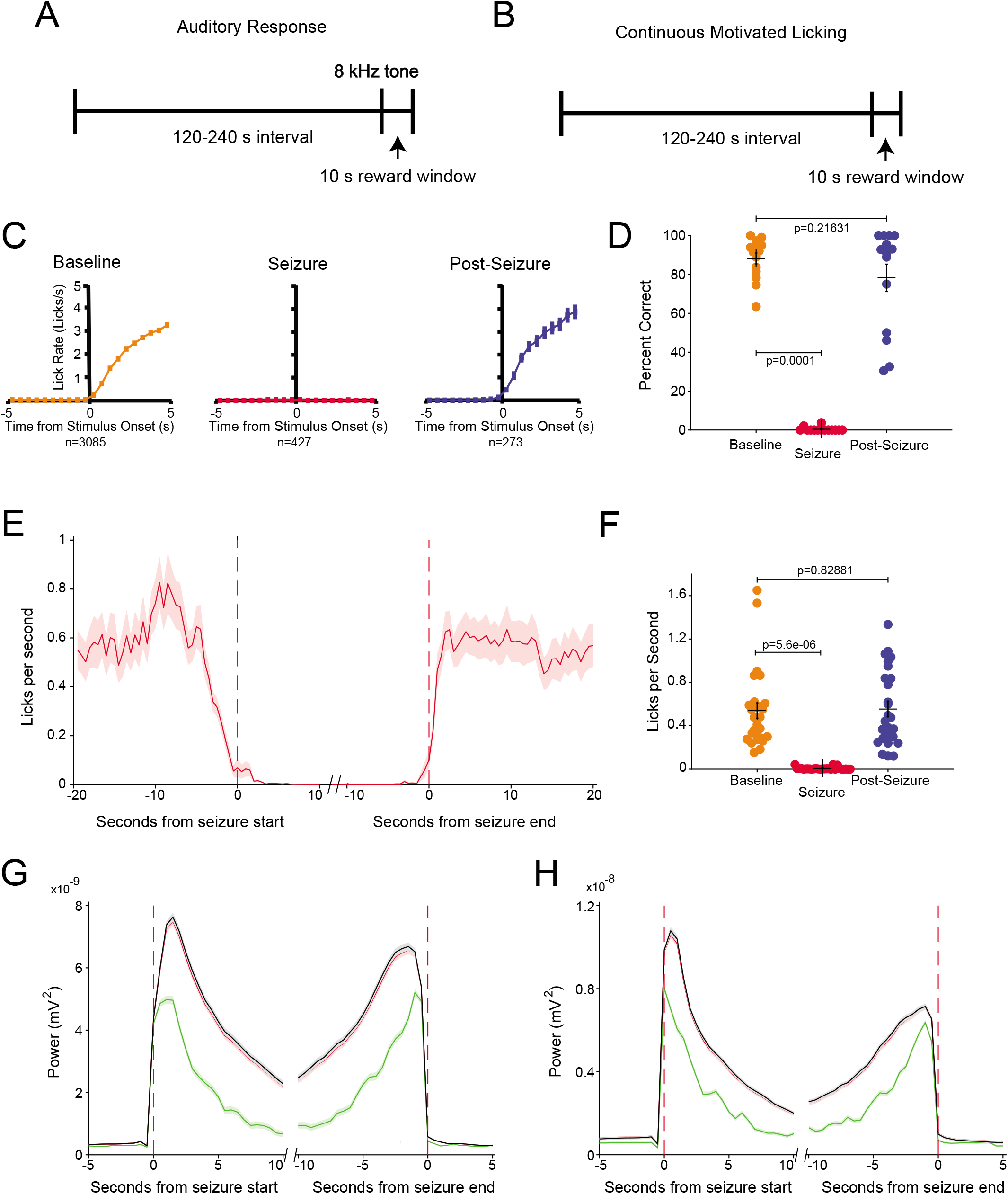
Behavior of GAERS around and during absence seizures. All measures of center are mean and error bars/regions denote standard error of the mean. **A:** design of conditioned auditory response task. Stimuli were presented at intervals of between 120 and 240 seconds (or when seizures were detected). An 8 kHz tone (intensity 45 dB) was used to signify the availability of a reward within a 10s window (n = 427 seizures from 14 animals for this task). **B:** design of continuous motivated licking paradigm. Rewards became available for 10s windows at intervals of between 120 and 240 seconds with no associated stimulus or signifer to encourage regular licking (n = 3146 seizures from 27 animals for this paradigm). **C:** lick rate following conditioned auditory stimuli delivered at baseline, during seizure, and in the 10 seconds immediately post-seizure. **D:** percent of conditioned stimuli responded to within the reward window and within the specified state for baseline, seizure, and immediate post-seizure periods for each animal showing decrease during seizure and recovery in post-seizure. **E:** dynamics of lick rate in 0.5s bins around seizure start and end times in continuous motivated licking paradigm. **F:** mean lick rates during baseline, seizure, and immediate post-seizure periods in continuous motivated licking paradigm showing decrease during seizure and recovery in post-seizure. **G:** dynamics of EEG power in the spike band (15-100 Hz) surrounding seizure start and end times for seizures with continued licking (green) and no licking (black). **H:** dynamics of EEG power in the wave band (5-9 Hz) around seizure start and end, comparing continued licking and no-licking seizures.

In the second task, designed to resemble the human RTT, the instructed repetitive tapping was approximated by providing unheralded sucrose rewards at varying intervals. This paradigm encouraged spontaneous licking at the spout at a mean rate of 0.54 ± 0.07 licks/second outside of SWDs (27 rats), and was therefore referred to as a sustained motivated licking task. We observed a decrease in lick rate from approx. 0.75 licks/second 10 seconds pre-SWD to a mean lick rate during seizures of 0.007 ± 0.002 licks/second (total of 3146 seizures), constituting a significant decrease from pre-seizure periods (p = 5.6e-6, Wilcoxon signed rank test). Within 2-3 seconds after seizure end there was an apparent recovery in lick rate, with a mean post-seizure rate of 0.55 ± 0.06 licks/second (p = 0.8288 relative to all non-seizure periods, Fig. 1F).

These results are to our knowledge the first demonstration of consistently impaired behavioral interactions with the environment during SWD in a rodent absence epilepsy model. This provides important face validity for the rodent model to investigate mechanisms of impaired behavioral interactions in human absence epilepsy. As further validation of the model, we sought to determine whether behavior might be spared in some SWD. In human absence epilepsy, behavior may be spared in some SWD especially in tasks that are less behaviorally demanding (Berman et al., 2010; Guo et al., 2016). In addition, in human absence epilepsy, the SWD showing spared behavior are significantly less physiologically severe based on magnitude and duration(Berman et al., 2010; Guo et al., 2016). Similarly, in the GAERS model we found that performance on the more demanding auditory response task was virtually always impaired during SWD (Fig 1d), whereas in the less demanding spontaneous licking task, approximately 5% of all SWDs (158/3146) demonstrated some persistent licking during SWD (Fig 1g,h). Again resembling human absence seizures, the SWD with spared behavior in the rodent model featured significantly lower EEG power in bands corresponding to both the wave (5-9 Hz, p = 3.213e-9, rank sum test) and the spike (15-100 Hz, p = 4.490e-10, rank sum test) components of the spike-wave oscillation. In addition, the mean duration of SWD with spared behavior (5.9 ± 0.6 s) was significantly shorter compared to SWD with impaired behavior (8.9 ± 0.2 s, p = 8.8e-9, rank sum test).

These behavioural results suggest that SWDs in GAERS have similar effects on consciousness as do absence seizures in humans. As such, neuronal activity during these SWDs may provide valuable insight into potential mechanisms of absence seizures and their symptoms.

### Total neuronal activity accompanying GAERS SWDs

Having established that GAERS SWDs were accompanied by impaired behaviour, we investigated the changes in neuronal activity that might cause these impairments. Our investigation of neuronal activity was targeted at representative cortical and thalamic regions known to be involved in SWD based on prior work. Cortical involvement in rodent SWD is most prominent in somatosensory cortex, particularly in the peri-facial areas (Meeren et al., 2002; Polack et al., 2007). To avoid potential extreme values in most intensely involved areas, we therefore recorded from somatosensory cortex but in the trunk region outside the face area. To investigate thalamic activity, we performed new analyses on previously acquired recordings of neuronal activity from ventral basal somatosensory thalamus (McCafferty et al., 2018). First, we studied the firing of 168 individually sorted neurons (Fig. 2A-D) in the somatosensory cortex around SWD initiation and during SWDs. The mean firing rate of these cortical neurons decreased from 3.7 ± 0.4 spikes/second during non-seizure to 2.6 ± 0.3 spikes/second during seizure periods (p=, 2.7e-9, Wilcoxon signed rank test). Interestingly, there was a transient peak in firing just at the point of seizure initiation, followed by a sustained decrease that lasted until seizure termination (Fig. 2E). A similar early peak in neuronal activity followed by sustained firing decreases was observed previously in thalamocortical neurons during SWD (McCafferty et al., 2018), and replicated in the present analysis of total thalamic neuronal firing (Fig. 2F). Analysis of the distribution of neuronal firing surrounding spike-and-wave complex (SWC) peaks (the most extreme voltage value of the spike) in the first second of seizures showed a higher oscillation frequency and higher firing rate than later times in seizures (Fig. 2G,H).

**Figure 2.**
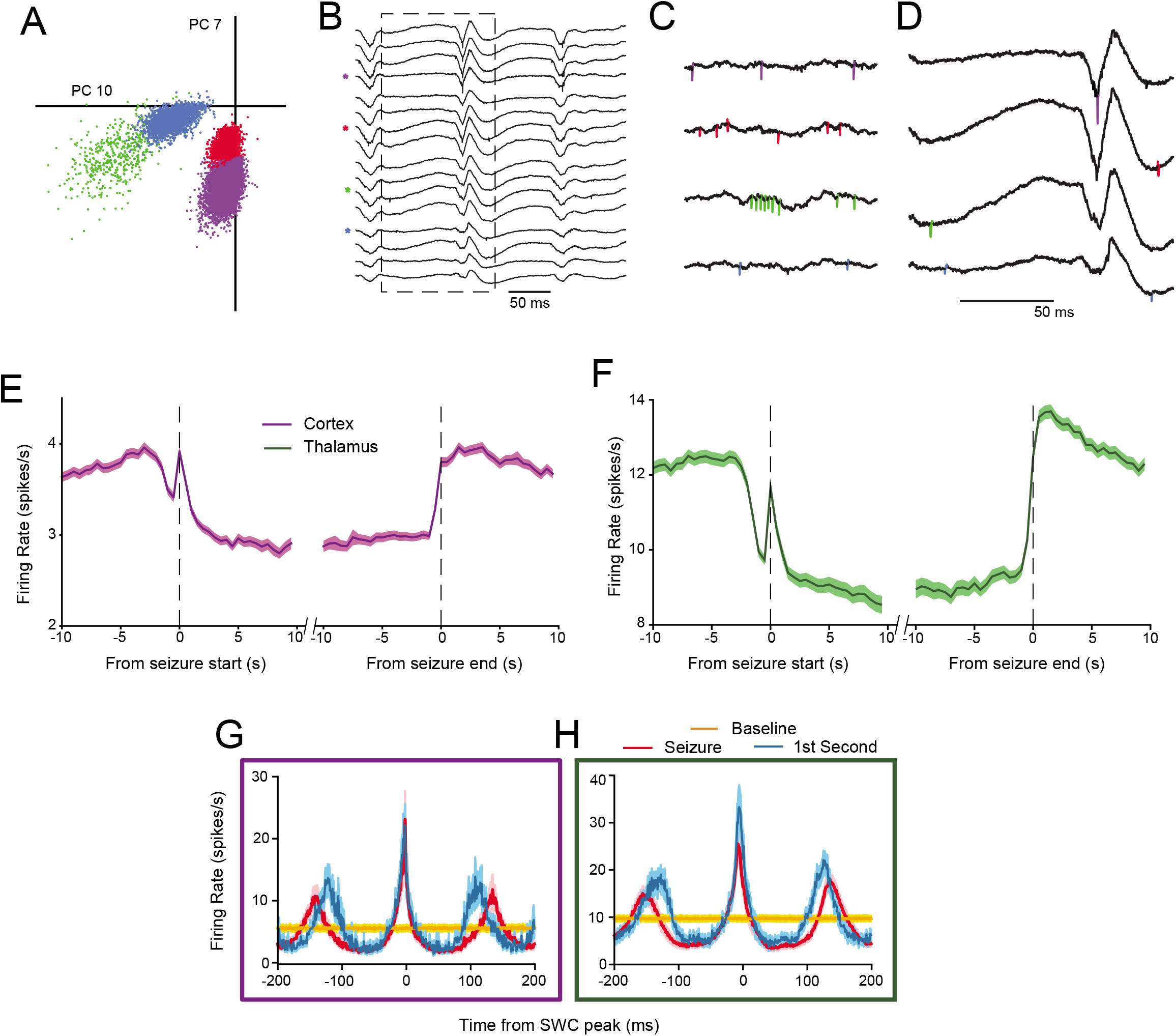
Total firing of cortical (n=165) and thalamic (n=164) neurons during absence seizures. All measures of center are mean and error bars/regions denote standard error of the mean. **A:** principle component values of waveforms of sample simultaneously-recorded cortical neurons. **B:** sample raw broadband (1.1 Hz to7.6 kHz) voltage traces from which neuron action potentials were extracted (channels of greatest action potential waveform amplitude indicated with asterisks). **C:** sample of firing of neurons from A and B during wakefulness without SWD. **D:** expanded view of mid-seizure section from B (dashed box in B) showing channels of greatest amplitude for each individual neuron. **E:** mean firing dynamics around seizure start and end of all cortical neurons in 0.5s bins, showing overall decrease in firing associated with seizure. **F:** mean firing dynamics around seizure start and end of all thalamic neurons (raw data from McCafferty et al., 2018) showing similar overall decrease. **G:** distribution of action potentials around spike-and-wave complex peaks in 1 ms bins of all cortical neurons, showing higher oscillation frequency of first second of seizure. **H:** distribution of action potentials around spike-and-wave complex peaks in 1 ms bins of all thalamic neurons, showing similar first second increased frequency.

### Diverse neuronal activity accompanying GAERS SWDs

Recent evidence suggests that cortical and thalamic neuronal activity during SWDs may not be as homogeneous as previously thought (McCafferty et al., 2018; Meyer et al., 2018). We investigated whether diversity of firing patterns existed in GAERS somatosensory cortical neurons, and found that the firing rate dynamics of these cells around SWD initiation tended to fall into one of four patterns (Fig. 3A). These were: a peak in firing at seizure initiation, followed by a return to baseline levels throughout (Onset Peak group (OP), 44 neurons, 28%), a sustained increase or decrease in firing throughout the seizure (Sustained Increase (SI) and Sustained Decrease (SD) groups, 15 and 59 neurons; 9% and 37% respectively), or no apparent change in firing rate associated with the seizure (No Change group (NC), 41 neurons, 26%). These patterns were distinctive and consistent within groups.(McCafferty et al., 2018) Interestingly, analysis of thalamic unit activity around SWD revealed that the same four patterns were apparent in these dynamics, with each neuron showing either an Onset Peak (18 neurons, 14%), Sustained Increase (4 neurons, 3%), Sustained Decrease (58 neurons, 46%), or No Change (45 neurons, 36%). Having established this diversity in mean firing rate we were also interested in whether the rhythmicity of cells differed according to the firing rate dynamic group (Fig. 3B). We found that the cortical Onset Peak and Sustained Increase groups had the most pronounced increases in rhythmicity of firing, with prominent peaks in firing during the spike phase leading to an overall increase in neuronal firing during SWD in these two neuron groups. In contrast, the Sustained Decrease and No Change neurons had less pronounced increases in rhythmicity (Fig 3B). In the Sustained Decrease neurons troughs were larger and longer than the peaks leading to an overall decrease in firing during SWD. In the No Change neurons the magnitude of peaks and troughs were relatively balanced, leading to little change in firing relative to baseline.

**Figure 3.**
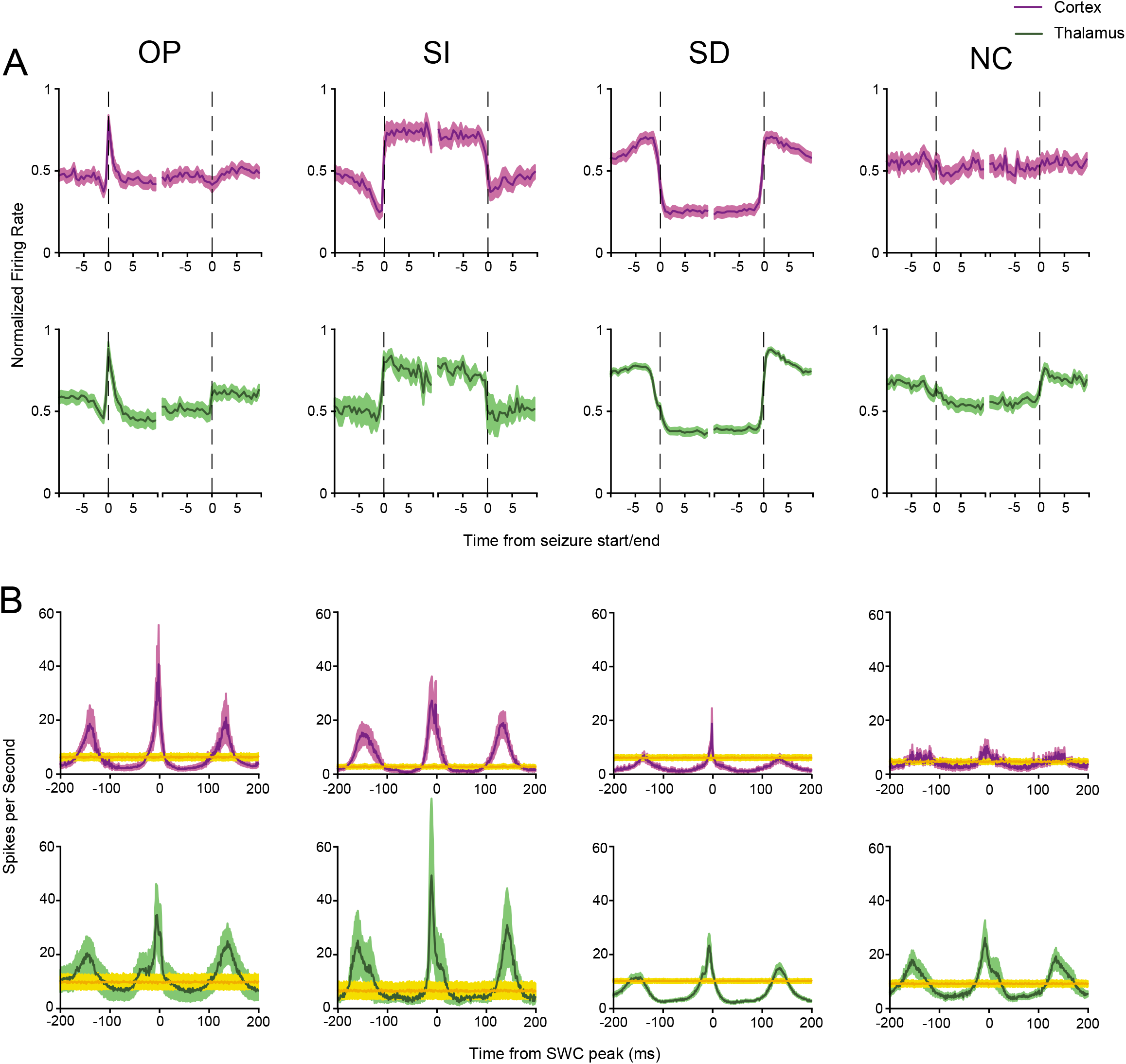
Firing rates and patterns of firing of subgroups of cortical and thalamic neurons during absence seizures. **A**: firing rate in 0.5 second bins (scaled relative to total range of firing in visualized period) for each group of neurons showing the distinctive dynamics around seizure onset/offset from which their names are derived: OP = onset peak, SI = sustained increase, SD = sustained decrease, NC = no change. **B**: mean distribution of action potentials in 1 ms bins around peaks of spike-wave cycles for the same groups. Inset values are measures of rhythmicity: total firing in the 50ms surrounding SWC peak divided by total firing in two 50 ms bins either side of the peak (−100:−50 ms and +50: +100 ms). Values expressed as mean ± standard error.

## Conclusions

These results are, to our knowledge, the first scientific demonstration of animal SWDs accompanied by an impairment of motivated behaviour similar to that observed in humans during absence seizures. This constitutes a valuable opportunity to investigate the neuronal mechanisms underlying the impairment – if particular groups of neurons and/or patterns of activity responsible for the loss of consciousness in seizure can be identified, then therapeutic interventions could target them to restore consciousness or even prevent seizures. We also show here the first evidence of specific and diverse patterns of neuronal activity accompanying these consciousness-abolishing SWDs, and suggest that investigating the characteristics and potential different roles of these groups of neurons may indicate prime targets for absence seizure therapeutics.

## Summary Methods

### Animals

Experiments were carried out under approval by the Yale University Office of Animal Research Support (OARS). All experiments were carried out with Genetic Absence Epilepsy Rats from Strasbourg (GAERS), an established polygenic rat model of absence seizures. Animals had access to food and water *ad libitum* unless otherwise noted, and were kept on a 12:12 hour light:dark cycle.

### Behaviour

All animals were implanted with fronto-parietal epidural screw electrodes under isoflurane anaesthesia for EEG recording and the identification of SWDs. Both behavioural paradigms were carried out in custom operant chambers (Med Associates Inc.). For the sensory tone detection task, rats were trained by increments to respond to an 8 kHz tone by licking at a port within 10 s in order to receive a reward bolus (90 μL) of sucrose water. For the sustained motivated licking paradigm, an unheralded reward bolus became available for 10 s periods at intervals varying from 150 to 210 s.

### Neuronal activity

Cortical data was collected using four-shank multi-electrode silicon probes (NeuroNexus) in the GAERS somatosensory cortex (coordinates from bregma AP – 3mm, ML ± 3 mm) and the OpenEphys digitization/acquisition system. Data was collected during 2-4 hour sessions during which rats were free to explore, rest, and seize in the recording chamber. Signals from each channel were band-pass filtered between 1.1 and 7603.8 Hz and digitized at 30 kHz and 192x gain. Thalamic data was used with permission from the dataset described in (McCafferty et al., 2018). Action potential spikes were extracted from the signal and clustered into separate neurons as previously described (McCafferty et al., 2018).

## Authors’ contributions

CMcC, HB, BG, PH, BS, FH and AD designed research and experiments; CMcC, BG, PH, BS and all other authors performed experiments and analyzed data, CMcC and HB wrote the manuscript with critical review by other authors.

